# Replicative cellular age distributions in compartmentalised tissues

**DOI:** 10.1101/305250

**Authors:** Marvin A. Böttcher, Benjamin Werner, David Dingli, Arne Traulsen

## Abstract

The cellular age distribution of hierarchically organized tissues can reveal important insights into the dynamics of cell differentiation and self-renewal and associated cancer risks. Here, we examine theoretically the effect of progenitor compartments with varying differentiation and self-renewal capacities on the resulting observable distributions of replicative cellular ages. We find that strongly amplifying progenitor compartments, i.e. compartments with high self-renewal capacities, substantially broaden the age distributions which become skewed towards younger cells with a long tail of few old cells. However, since mutations predominantly accumulate during cell division, a few old cells may considerably increase cancer risk. In contrast, if tissues are organised into many downstream compartments with low self-renewal capacity, the shape of the replicative cell distributions in more differentiated compartments are dominated by stem cell dynamics with little added variation. In the limiting case of a strict binary differentiation tree without self-renewal, the shape of the output distribution becomes indistinguishable from the shape of the input distribution. Our results suggest that a comparison of cellular age distributions between healthy and cancerous tissues may inform about dynamical changes within the hierarchical tissue structure, i.e. an acquired increased self-renewal capacity in certain tumours.

## 1. Introduction

Many tissues in multicellular organisms resemble a compartmentalised structure with a hierarchy of cells at different stages of differentiation and function. This hierarchy is usually fuelled by a few stem cells that ideally can self-renew indefinitely, whereas the majority of the tissue consists of shorter-lived differentiated cells that emerge from these stem cells [1-3].

In most tissues it is thought that stem cells divide infrequently, while their progenitors and further differentiated cells divide more frequently to ensure tissue function under homeostasis [4]. Such structures allow both the production of many cells in a short time and the reduction of the risk for the accumulation of somatic mutations within the stem cell compartment [1, 5-10].

Due to these pronounced dynamical disparities in hierarchical tissues, replicative age — the number of divisions a cell has undergone — can be an important observable providing information about the structure and cellular dynamics within these tissues. Since many somatic mutations are acquired during cell divisions [11, 12], we would expect replicative age also to be strongly correlated with different cancer risks in different hierarchical tissues [13-15]. In the context of ageing, the focus is typically on changes within the stem cell compartment, since stem cells have the ability to self-renew and persist long enough to become relevant for organismal ageing [16, 17]. It is generally assumed that replicative cell age in downstream compartments is a good proxy for replicative stem cell age. For example, some of us previously developed and tested a mathematical model for human hematopoietic stem cell ageing based on replicative ages in human lymphocytes and granulocytes [18].

Cellular dynamics in hierarchically organised tissue structures can be hard to explore experimentally due to the large scaling difference between differentiation levels [19] and the challenges to correctly assign cells to specific differentiation stages. One possibility to assess the age distribution of a cell population is to measure the telomere length of the cellular chromosomes. Telomeres are the protective, non-coding ends of chromosomes, consisting of the same short DNA sequence repeated thousands of times. Telomeres typically shorten with each cell division [20-22]. Cells with critically short telomeres enter replicative senescence, which is thought to be a cancer suppression mechanism [23]. Moreover, critically short telomeres are often associated with genome instability and corresponding increased risk of cancer [24]. For our purpose, telomere length distributions can be thought of as a measure for the cellular replicative age distribution. They can be assessed in tissue samples [25, 26] which are for example especially accessible in differentiated tissue in the hematopoietic system and thus in principle would also allow for some time resolution within healthy human individuals [18].

However, to truly infer stem cell dynamics from replicative age distributions in differentiated cells, a more detailed understanding of the differentiation processes and their impact on the replicative age distributions is necessary. Here, we develop a mathematical framework that allows us to describe the distribution of replicative cellular ages across several hierarchical levels of differentiation. Thereby, we demonstrate how the distribution of replicative ages in differentiated cell populations can provide insights into the properties of the dynamics within the underlying tissue.

## 2. Model

In the following, we present a mathematical description for the replicative age distributions within compartmentalized tissue structures (fig. 1). First, we discuss the simplest case of only two compartments - one stem cell compartment and the focal downstream progenitor compartment. We then ask what is the distribution of replicative ages of cells in the progenitor compartment provided a continuous influx of cells from the stem cell compartment. We do not discuss the time dynamics on the stem cell level explicitly. The time evolution of replicative age distributions in stem cell compartments and the resulting potential influx distributions for progenitor compartments are discussed in detail in [18].

**Figure 1:**
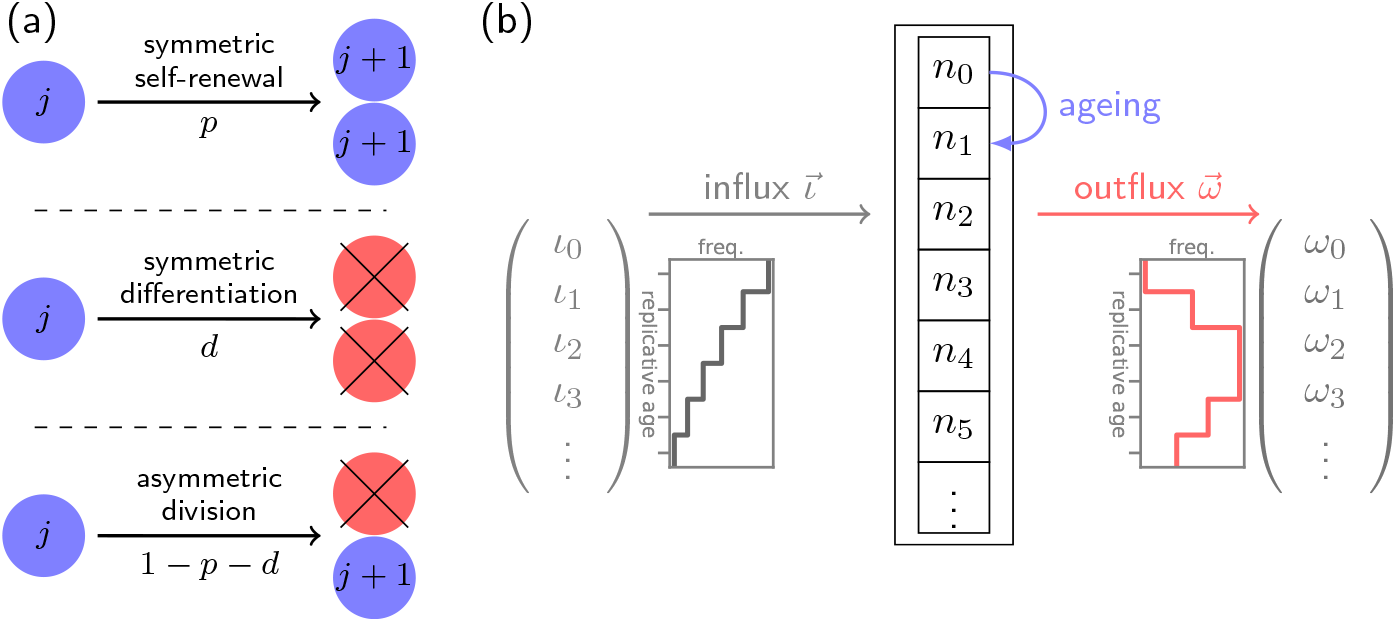
Sketch of the basic model. (a) Three different modes of cell division in the focal progenitor compartment. Blue cells are cells within the compartment, red cells differentiated and leave the compartment. The replicative age of a cell in the specific compartment is *j*, increasing by one in each cell division. (b) Fill model for ageing in progenitor compartment. The number of cell in each age class is *n_j_*, and all cells age according to the modes of cell division (a). The cell influx 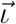 into the compartment has a certain distribution of replicative age *ı*_*j*_. The cell outflux from the compartment includes all differentiating cells and is denoted by the distribution 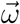.

We assume that in the progenitor compartment there are *n_j_* cells of each replicative age class *j*. Progenitor cells divide with proliferation rate r and after each division the replicative age of both daughter cells increases by one *j* → *j* + 1. Each daughter cell can in principle take a different cell fate that contributes differently to the distribution of replicative ages (fig. 1A). In general, the following outcomes are possible after a single cell division

i. With probability *p* a cell self-renews symmetrically, both daughter cells stay in the same compartment and increase their cellular age by one (*n_j_* → *n_j_* − 1, *n*_*j*+1_ → *n*_*j*+1_ + 2)
ii. With probability *d* a cell differentiates symmetrically, effectively removing it from the compartment of differentiated cells (*n*_*j*_ → *n*_*j*_ − 1).
iii. With probability 1 − *p* − *d* a cells divides asymmetrically, with one cell staying in the pool of differentiated cells while the other cell leaves the compartment [27] (*n_j_* → *n_j_* − 1, *n*_*j*+1_ → *n*_*j*+1_ + 1).

We choose the influx of cells from the stem cell compartment to be a constant rate *ι*_*j*_ that might differ for each cellular age *j*. Below we will give explicit examples for different distributions of *ι*_*j*_. We assume the dynamics on the stem cell level to be much slower compared to downstream compartments and hence consider the influx *ι*_*j*_ into the progenitor compartment to be constant over time.

Using the above, we can formulate differential equations for the change of the number of cells in each age class *n*_*j*_. Thereby, we account for the loss of cells due to proliferation and subsequent differentiation and gain of cells due to symmetric self-renewal and cell influx from the stem cell compartment,

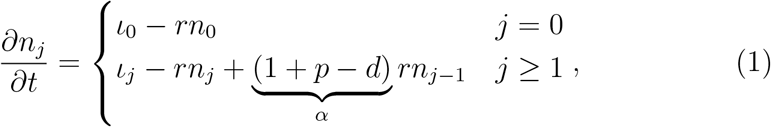

where we set *α* = 1 + *p* − *d* to be the self-renewal parameter which critically determines the most relevant results of our model. Since *p* and *d* are probabilities with *p* + *d* ≤ 1, the self-renewal parameter can be in the range 0 ≤ *α* ≤ 2. However, since we are interested in homeostasis and not an exponentially growing tissue, the symmetric division probability *p* in our case must be smaller than the symmetric differentiation probability *d* and therefore 0 ≤ *α* ≤ 1.

The above system of ordinary differential equations can be solved analytically (see appendix E). However, as we assume that the dynamics on the level of stem cells is much slower compared to progenitor compartments we can investigate the equilibrium solutions 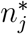 to equation 1 for each age class *j*. The equilibrium solutions can be obtained via recursion by setting 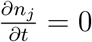 (see appendix A). The general solution is

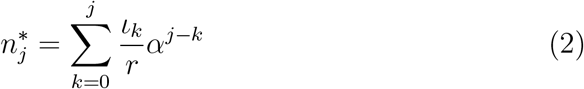

which is equivalent to a convolution sum of the influx *ı*_*k*_ and 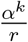 between zero and *j*.

**Figure 2:**
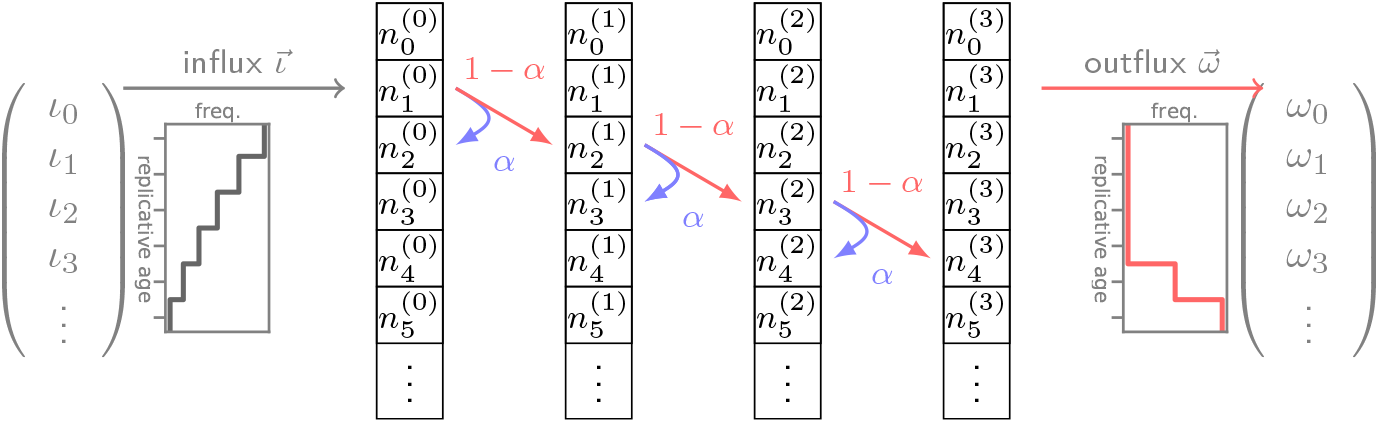
Several downstream compartments amplifying the rate of cell production from influx to outflux. In each compartment there are self-renewal or differentiation processes as described in figure 1. Each cell division thereby leads to an increase of replicative age and changes the age distribution of the corresponding compartment. Self renewal occurs proportional to the self-renewal parameter *α*, whereas differentiated cells are produced with 1 − *α* and go into the next downstream compartment. The compartment number *c* is shown as superscript, the total number of compartments is *C* = 4.

### 2.1. Multiple compartments

In reality, most tissues will consist of multiple progenitor cell compartments.

It is thus natural to ask how multiple downstream compartments affect cellular age distributions. To answer this question, we can generalise our previous framework. Differentiated cells in a downstream compartment are either produced by symmetric differentiation with probability *d* or by asymmetric division with probability 1 − *p* − *d*. If we denote the output of cells per unit of time for each age class as *ω*_*j*_, we can write

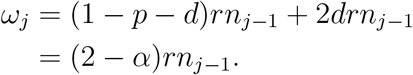

To allow for multiple compartments, we can identify the output distribution of a compartment *c* and the input distribution of the next downstream compartment *c* + 1,

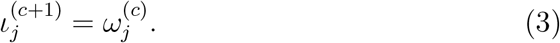

#### Total cell outflux

For our purpose it is desirable to compare the effect of different tissue structures, that is a different number of total compartments *C*, but with the same tissue function, that is the same total outflux of fully differentiated cells. In our model, the total outflux of differentiated cells Ω = ∑_*j*_ *ω*_*j*_ is determined by the total influx of cells *I* = ∑_*j*_ *ı*_*j*_, the number of compartments *C* and the self-renewal parameter *α*. We therefore choose *α* such that the total output of cells remains constant, i.e. assuring certain replenishing needs of a specific tissue. For this we formulate differential equations for the change of the total number of cells 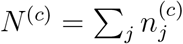 in each of the compartments *c* with a compartment specific proliferation rate for each cell *r*^(*c*)^ by collecting all influx and outflux terms:

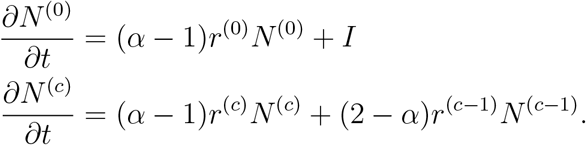

Here, *I* is the total influx into the first compartment (*c* = 0) (i.e. the sum of all direct stem cell derived progenitors per time unit). The total outflux Ω is related to the number of cells in the last compartment *N*^(*C*−1)^ via

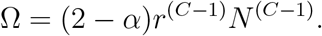

Under steady state conditions, the above equations can be solved explicitly for the self-renewal parameter *α* (see appendix B):

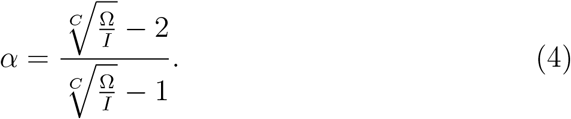

This allows us to adjust the self-renewal parameter *α* such that the outflux Ω remains constant given an influx *I* for any number of compartments *C*. However, since the self-renewal parameter is constrained 0 ≤ *α* ≤ 1 (see above section) the minimum amplification of cell production is given by 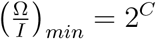 corresponding to *α* = 0.

### 2.2. Properties of the replicative age distribution

#### Mean and variance

The mean and variance of the replicative age distribution under steady state conditions can be calculated analytically, see appendix C. The mean *μ* of the replicative age distribution in the progenitor compartment increases compared to the influx based on the self-renewal *α* to

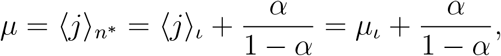

where 〈*j*〉_*n*\*_ is the first moment of the replicative age distribution in the focal progenitor compartment and 〈*j*〉_*ı*_ = *μ*_*ı*_ is the average replicative age of the influx. Note that the average replicative age of the outflux *μ*_*ω*_ = 〈*j*〉_*ω*_ is increased by one to account for the extra differentiation step

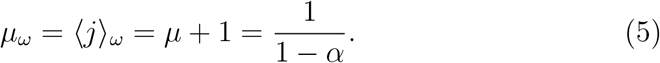

The minimal increase of the mean between influx and outflux for no self-renewal (*α* = 0) is therefore equal to one.

The variance *σ*^2^ of the replicative age distribution increases similarly as the mean above

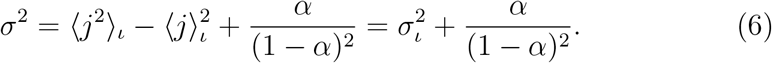

Here, 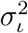 denotes the variance of the replicative age distribution of the influx.

Generally, also the higher moments 〈*j*^*γ*^〉_*n*\*_ of the replicative age distribution can be calculated based on the moments of the influx distribution 〈*j*^*β*^〉_*ı*_ with *β ≤ γ*. The corresponding calculations and results are shown in the appendix section C.

#### Limiting behavior

For very low self-renewal, *α* ≪ 1, the only age class of influx that significantly contributes to the age distribution 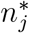 in equation 2 is *ι*_*k=j*_, as it is in zeroth order of *α*. The influx of all other age classes is of higher order of self-renewal *α* and will therefore vanish for *α* ≪ 1 such that

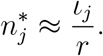

Hence, the outflux distribution will look approximately like the influx distribution.

To evaluate the impact of the progenitor compartment on the replicative age distribution in the limit of high self-renewal 1 − *α* ≪ 1, we rewrite equation 2 to

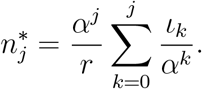

The limiting behavior therefore strongly depends on the age distribution of the influx *ι*_*k*_. If the influx has an upper bound *K* on replicative age, such that for all *k* ≥ *K* holds *ı*_*k*_ ≪ *α*^*k*^, the sum in the above equation is constant and the distribution of replicative age will decline exponentially

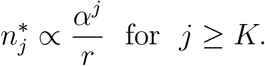

If, on the other hand, the influx distribution *ι*_*k*_ is not declining fast enough and is in the same order as *α*^*k*^ (*ι*_*k*_ ≥ *α*^*k*^), we can not make a general prediction for this limit.

## 3. Results

It seems natural to suspect that the specific distribution of replicative ages in downstream compartments strongly depends on the distribution of cellular ages within the stem cell compartment. In the following, we present the resulting age distributions for various different influx distributions. Additionally we will compare tissue structures with many subsequent downstream compartments and a low probability for self-renewal against having only very few compartments with a high probability for self-renewal.

An important parameter for the age distribution in the progenitor compartment is *α* = 1 + *p* − *d* which depends on both the probability for symmetric splitting and symmetric differentiation and critically defines the total size of the compartment as well as the amount of cells produced (appendix B). For a compartment model of hematopoiesis with many differentiation steps as for example in [1, 28], *α* would be around 0.3, whereas for other models with fewer compartments *α* would need to be higher to allow for sufficient output of fully differentiated cells per unit time [29-31].

### 3.1. A single progenitor compartment

Here we discuss the distributions of replicative age in the special case of a single progenitor compartment given four different influx distributions from the stem cell compartment. All distributions are calculated analytically and the corresponding calculations can be found in the appendix D.1 and 3.1.3. Realisations of the resulting replicative age distributions are shown in figure 3.

**Figure 3:**
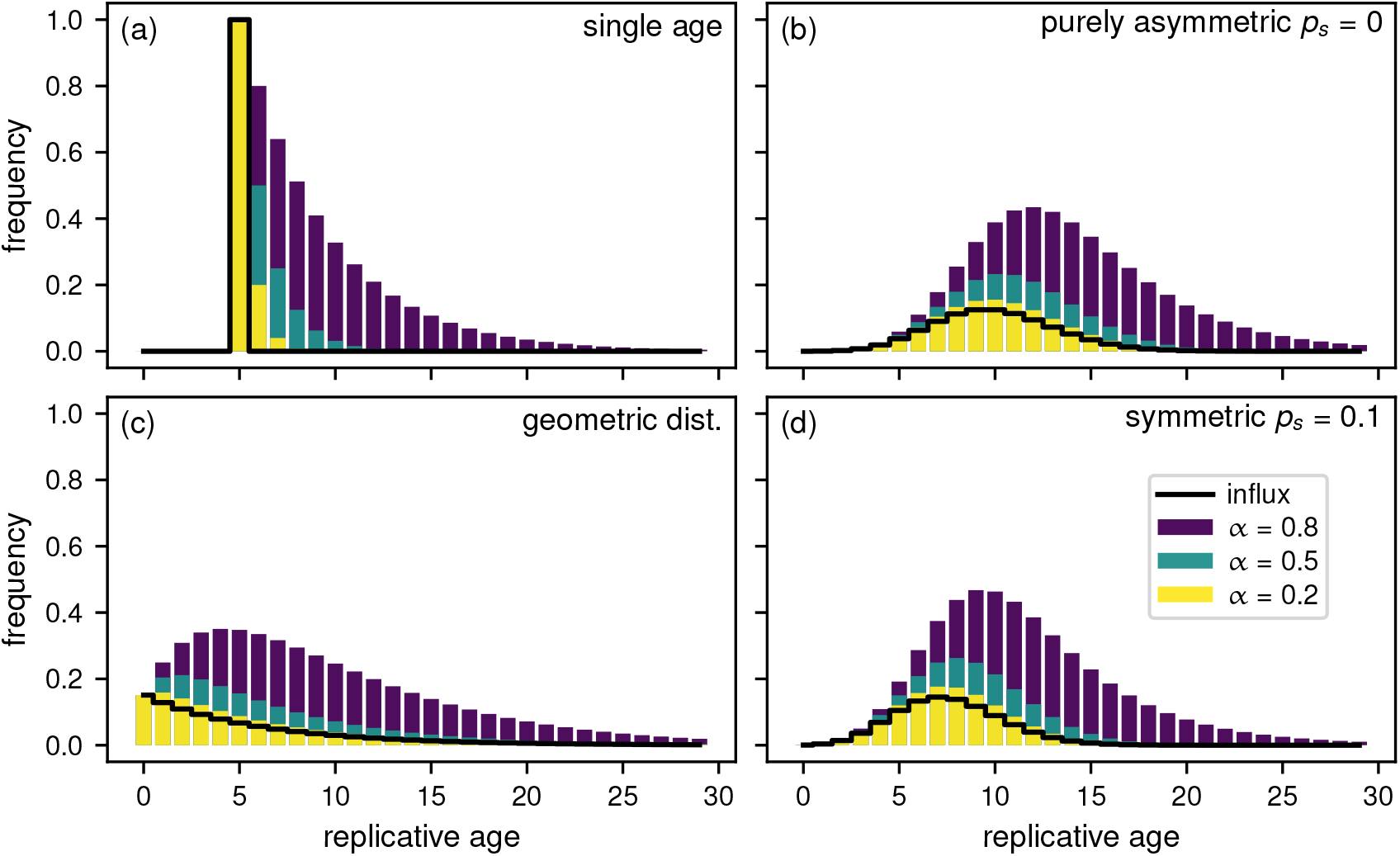
Distributions of replicative age in the first progenitor compartment for varying influx distributions from the stem cell compartment. (a) Influx of only a single replicative age *ı_k_* = *δ*(*k* − *v*) with parameter *v* = 5. (b) Influx given by a geometric distribution with many young and fewer old cells *ı_k_* = *λ*^*k*^(1 − *λ*). The distribution parameter is *λ* = 0.85. (c) Model based influx for purely asymmetric divisions on the stem cell level resulting in a Poisson distribution 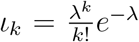 [18]. Parameter *λ* = 10. (d) A model based influx with symmetric divisions (probability *p_s_* = 0.1) also result in differently normalized Poisson distribution 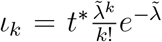 with a more pronounced difference to the age distribution for purely asymmetric divisions at older ages of the stem cell pool. In (c) and (d) the underlying parameters for *λ* and 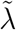 are the same (section 3.1.3 for details).

#### 3.1.1. Identical replicative cellular age influx

We first discuss the simplest case for a cellular age distribution on the stem cell level that is all stem cells have identical replicative age *v*. This results in a delta function input *ı_k_* = *r_s_δ*(*k* − *v*), where *δ*(*x*) is the Dirac delta distribution and *r*_*s*_ is the rate of cell production. Together with equation 2 this implies for the age distribution

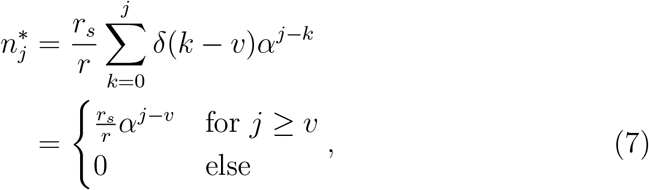

The resulting distribution is shown in figure 3A. Cellular ages within the single progenitor compartment follow an exponential distribution that approaches zero faster for smaller self-renewal parameters *α* and always has the maximum at the influx replicative age *v*.

#### 3.1.2. Geometrically distributed replicative cellular age influx

The former section is of course an oversimplification. We expect some form of distributed cellular ages on the stem cell level. We first discuss the possibility of a geometrically distributed replicative age *ı_k_* = *r_s_λ*^*k*^(1 − *λ*) with distribution parameter *λ* and total cell influx *r*_*s*_ as input from the stem cell level. This replicative age distribution resembles the distribution in the first progenitor compartment for an influx with identical replicative age from the stem cell compartment as shown in the previous section (Sec. 3.1.1); it would therefore correspond to the second downstream compartment for that specific influx.

The resulting age distribution within this progenitor compartment - equation 2 - can be solved analytically (see appendix D.1):

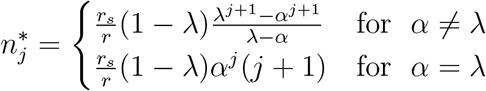

These age distributions are shown for different self-renewal parameters α in figure 3B. For low self-renewal, the shape of the replicative age in the progenitor compartment strongly resembles that of the influx distribution, i.e. a monotonically decreasing function of replicative age. For large self-renewal *α* ≥ 0.5, however, the distribution of replicative cellular ages in equilibrium becomes increasingly independent of the influx distribution and very similar to the age distributions resulting from other influx distributions, see e.g. figure 3 C and D.

#### 3.1.3. Influx from stem cell pool with random stem cell divisions

We previously investigated the dynamics within the stem cell compartment given that stem cell proliferations are independent and distribution times are exponentially distributed [18]. Once a stem cell is picked for division it either divides symmetrically with probability *p_s_*, resulting in two stem cells, or asymmetrically with probability 1 − *p*_*s*_, resulting in one progenitor and one stem cell. Now we ask how influx from such a stem cell pool percolates through the hierarchy.

##### Asymmetric stem cell divisions

Exclusively asymmetric divisions (*p*_*s*_ = 0) on the stem cell level result in a Poisson distribution of replicative age [18] and the corresponding influx into the progenitor compartment is given by

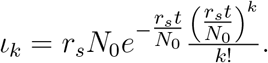

The distribution depends on age *t*, proliferation rate *r*_*s*_, as well as the initial number of cells *N*_0_ in the stem cell compartment. We can set 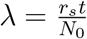 to see that this is a Poisson distribution multiplied by *r_s_N*_0_:

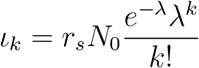

with a time dependent rate parameter *λ* = *λ*(*t*).

The corresponding sum from equation 2 can be solved analytically (see appendix D.2) and the distribution of replicative age becomes

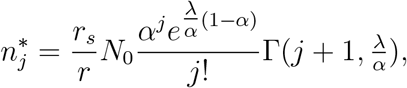

where 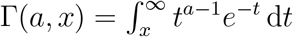 is the upper incomplete gamma function [32].

The above distribution of replicative age is shown in figure 3C for various values of the self-renewal probability *α*. The normalization factor *r_s_N*_0_ is set to one, as this does not change the general shape of the underlying distribution. Similar to our previous observations, the age distribution is heavily skewed towards younger cells. This effect is more pronounced for higher values of *α*, corresponding to more cells in the compartment.

##### Symmetric stem cell divisions

The age distributions for a growing stem cell compartment due to occasional symmetric stem cell self-renewals with probability *p*_*s*_ > 0 is given by [18]

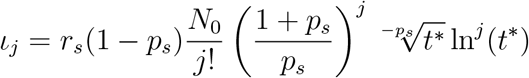

with 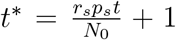. Here the exact distribution depends explicitly on the initial number of stem cells *N*_0_ and the ageing factor *t*\*, which itself depends on the relative increase of the stem cell pool size during time *t* given a symmetric division probability *p*_*s*_ and a proliferation rate *r*_*s*_. However, the distribution is again a Poisson distribution with a different normalization. This becomes apparent if we substitute 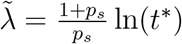 and get

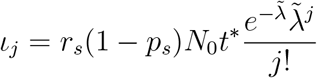

The solution of the convolution sum in equation 2 is therefore the same as for purely asymmetric stem cell divisions and the corresponding calculations are identical (if we exchange 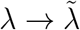) (appendix 3.1.3),

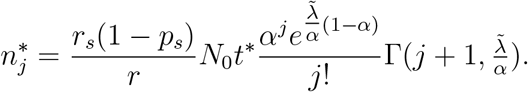

The shape of the resulting influx distribution therefore varies only slightly from the asymmetric case and differences in the age distribution of the progenitor compartment are minimal (fig. 3 C and D). However, the difference in average replicative age on the stem cell level is conserved in the progenitor compartment and still can be used to distinguish between those processes on the stem cell level [18].

### 3.2. Multiple compartments

In most organs, the maturation of functional tissue specific cells requires multiple stages of differentiation. We therefore generalize our approach above and discuss the impact of multiple subsequent non-stem cell compartments on the replicative age distribution within such hierarchical tissue organizations.

#### Impact of the number of compartments

In order to deduce the impact of the number of compartments on the age distributions, we vary the number of compartments by simultaneously keeping the final outflux of cells constant. This requires an adjustment of the self-renewal parameter *α* accordingly and is motivated by the idea that certain tissues might require a certain constant cell replenishment per unit time, but this could in principal be achieved in different tissue architectures. We use the same principal influx distributions from the stem cell compartment discussed above, see figure 3. Solutions in this section were obtained by numerically calculating the sums of equation 2.

Figure 4 shows the resulting replicative age distributions for a broad range of compartment numbers. Interestingly, the age distribution in the final compartment is very sensitive to the number of compartments, even though the total cell number amplification of the compartments is the same by construction. For a large number of compartments and corresponding small self-renewal a the shape of the influx distribution is basically conserved through all stages of the hierarchy, especially for the extreme case of a purely binary tree (*α* = 0) where the shape of the distributions is unchanged but only shifts towards older replicative age.

**Figure 4:**
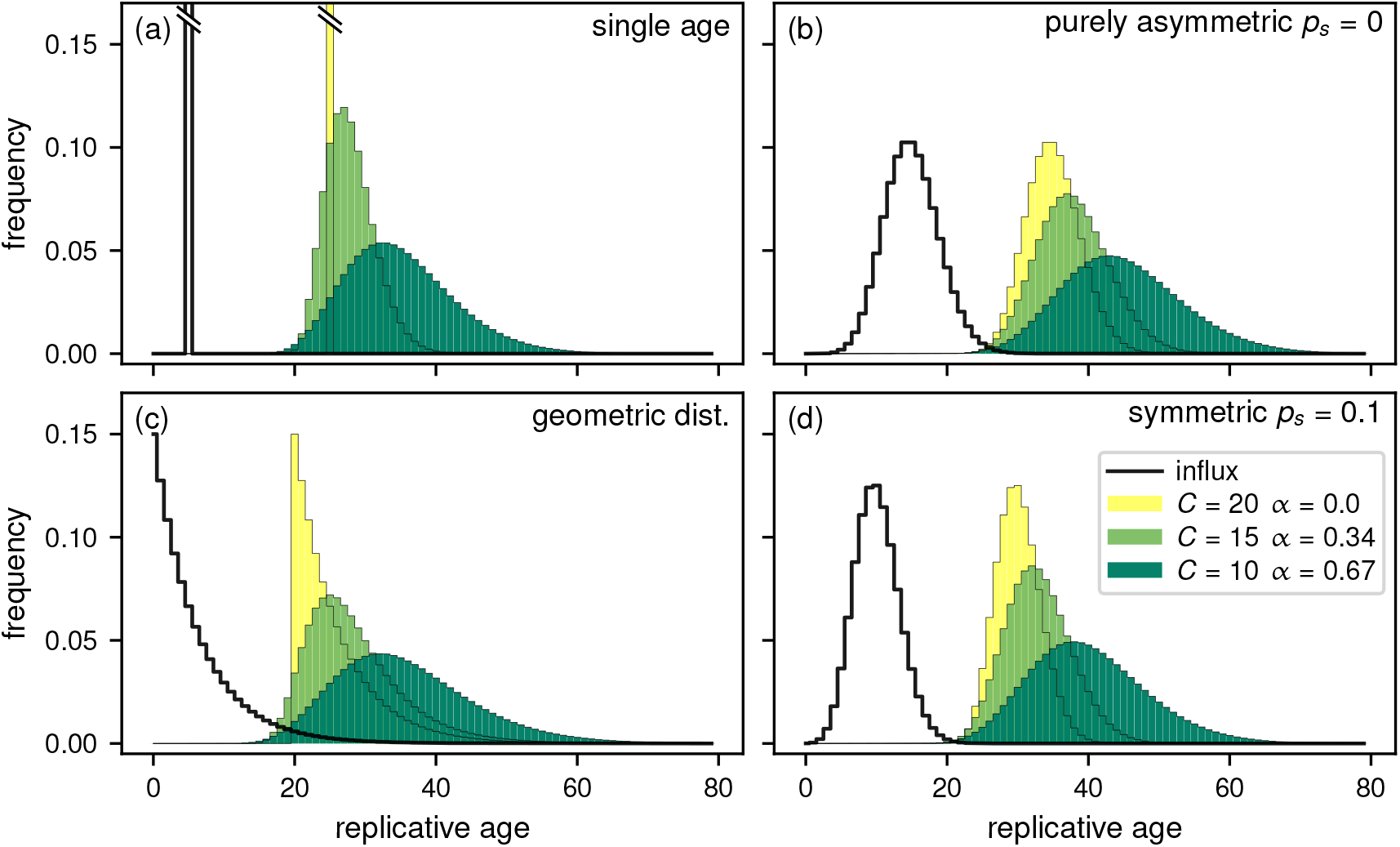
Comparison of different total number of progenitor compartments *C* for different influx age distributions. The self-renewal parameter *α* is adjusted such the total outflux Ω is the same for each *C*. The influx distributions are the same as in figure 3: (a) Influx with a single age. (b) Influx age geometrically distributed. (c) Model based influx for purely asymmetric divisions on the stem cell level. (d) Model based influx with symmetric divisions. For comparison of the influx 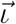 with the resulting outflux 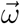, the distributions are normalized (all parameters as in figure 3).

For the other extreme case of only one or two downstream compartments (*α* ≈ 1), the distribution of replicative age is almost flat, such that the frequency of young cells is the same as the frequency of very old cells. Note, however, that in this case the steady state assumption might be violated as the time to reach homeostasis, i.e. the state where the system does not change anymore, might exceed realistic biological time scales. This is shown in the appendix section E in more detail.

However, distributions of replicative age become similar already for intermediate, but biologically still high, values of self-renewal *α* ≈ 0.5. It might therefore be impossible to distinguish between age distributions on the stem cell (influx) level from measurement in the differentiated tissue alone, provided there is considerable self-renewal in non-stem cell compartments. This is especially surprising considering the extreme differences in influx distributions, for example delta distributed (fig. 4A) and Poisson distributed (fig. 4C and 4D), which become seemingly undistinguishable in downstream compartments (at equilibrium). This effect is reminiscent of the law of large numbers for random variables, where the sum of independent random variables tends to a normal distribution regardless the actual distribution of the random variable.

#### Mean and variance through multiple compartments

In a system with multiple downstream compartments it is also interesting to see how mean and variance of replicative age change from compartment to compartment. As shown in equations 5 and 6 for a single progenitor compartment, mean and variance increase linearly from compartment to compartment with a slope of 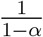 for the mean and 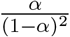 for the variance. Strong self-renewal therefore has a more pronounced effect on the variance than the mean due to the quadratic term in the denominator.

Figure 5 shows the mean and variance of replicative age for multiple subsequent compartments for different total number of compartments, but as above with the same tissue function, that is the same overall cell production. In this example, the variance for strong self-renewal, *α* = 0.67, at the second out of ten compartments is already larger than in the last compartment for the case of lower self-renewal, *α* = 0.34, even though there are five compartments more in the latter case. The impact on the mean of the distribution throughout the compartments is not nearly as pronounced. Since both, mean and variance, only depend on self-renewal *α* and the number of compartments, in principle stem cell dynamics can be inferred from comparing mean and or variance of telomere length distributions over time [18, 33], as long as the general tissue structure and dynamics does not change.

**Figure 5:**
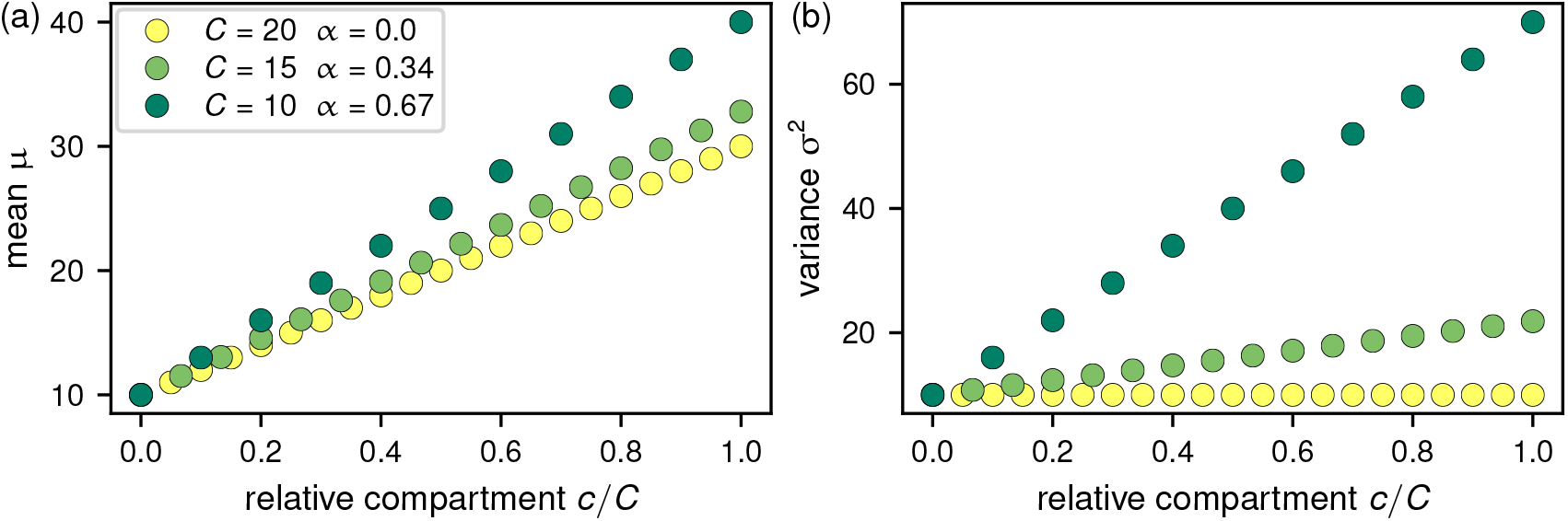
Mean *μ* and variance *σ*^2^ of replicative age distributions per compartment. Influx age distribution is the Poisson distribution with a mean and variance of *λ* = 10 as in figure 3(c); the x-axis shows the progression through the compartments 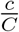. To compare different tissue structures, the self-renewal parameter *α* is adjusted for the same output of cells as in figure 2. (a) The mean of replicative age increases slightly faster for high self-renewal. (b) The variance of replicative age increases also linearly, however, the impact of the self-renewal parameter *α* is much more pronounced: for *α* = 0 there is no change, but for *α* = 0.67 there is a drastic increase of the standard deviation per compartment.

#### Change of replicative age distribution in CML

Chronic myeloid leukemia (CML) is a cancer of the hematopoietic system that can be characterized by enhanced self-renewal of cancerous cells in the progenitor compartments compared to healthy cells [34]. Here we compare the replicative age distribution for different self-renewal probabilities in the same tissue structure. For this, the tissue consists of 29 downstream compartments with either self-renewal probability *p* = 0.15 for healthy cell [1] or self-renewal *p* = 0.28 for cancerous cells [34] and without asymmetric division (*d* = 1 − *p*), leading to self-renewal parameters *α*_healthy_ = 0.3 or *α*_CML_ = 0.56.

The resulting distributions are shown in figure 6. For CML both mean and standard deviation are strongly increased compared to healthy hematopoiesis, which can be measured by comparing telomere length distributions during treatment of the disease [35]. We accordingly expect that both mean and standard deviation will decrease under successful treatment, when selfrenewal in progenitor compartments normalizes again, which is consistent with available clinical data [36].

**Figure 6:**
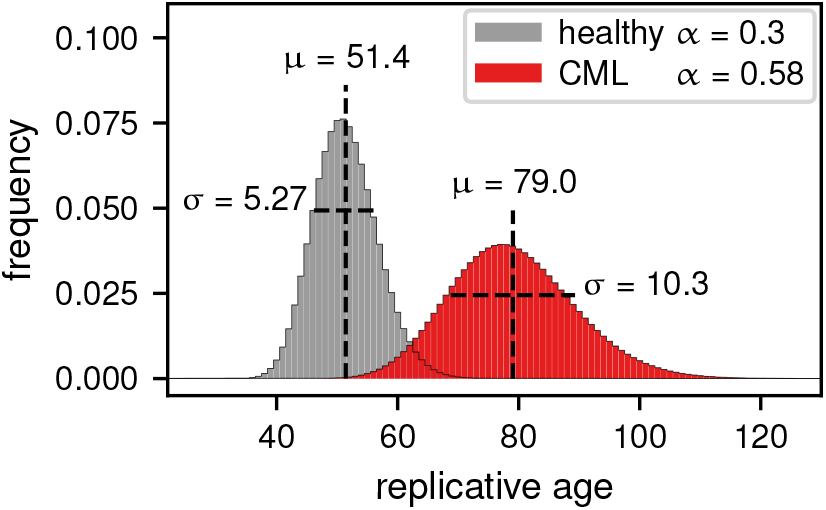
Replicative age distributions for healthy hematopoiesis and for hematopoiesis under chronic myeloid leukemia (CML). The self-renewal parameter *α* is the same in all 29 downstream compartments, *α* = 0.3 for healthy and *α* = 0.56 for cancerous hematopoiesis [34]. With CML, the replicative age distribution is much wider and shifted to a higher mean.

## 4. Discussion

While the age structure of cells within a tissue is driven by the age structure of the tissue specific stem cells, the progenitor compartments can substantially alter this age distribution. From a perspective of signal processing, they act as a filter that transforms an input signal (in our case a distribution) into an output signal. The properties of this filter are restricted by the biological structure of the tissue. Two limiting cases are of particular interest:

i. Focussing on a compartment that is weakly amplifying (*α* ≪ 1), such that the number of output cells is approximately twice the number of input cells, the replicative age distribution in the progenitor compartment resembles that of the influx distribution. Only the average age of the cells is then increasing with the compartment number, even in tissues with many subsequent downstream compartments, such as blood [1].
ii. For intermediate to high self renewal (large *α*), the distributions of cell replicative age in a differentiated tissue with multiple progenitor compartments are virtually indistinguishable from one another, even for influxes with completely different replicative age distributions. Measuring replicative age distributions in differentiated tissue, for example via telomere lengths [18, 33], may therefore be more informative about the tissue structure and dynamics than the dynamics within the long lasting stem cell level.

Cellular age is explored in many experimental studies (e.g. [37] gives a nice overview) and in multiple models [38, 39]. Some of these models also take the effect of replicative ageing into account [18, 39, 40]. However, replicative ageing in differentiated tissues is often overlooked, since here the cell turnover is very high and mutation accumulation as well as loss of function in these cells might not be as clinically relevant as in stem cells or early progenitor cells. On the other hand, we show that understanding the replicative ageing of differentiated cells and the resulting age distributions in the cell population can lead to a much better understanding of tissue dynamics from measurements. Previous models of replicative ageing in a tissue hierarchy including stem cells and progenitor cells focussed strongly on the total replication limit of cells [41, 42]. However, it is not clear whether or not fully differentiated cells are close to the end of their replicative life-span in vivo. Thus, we instead addressed the change in replicative age distributions going through possibly multiple rounds of differentiation without taking a potential maximum replicative age into account.

When comparing the distributions of replicative age between individuals or at different time points (or, for most practical purposes, their average and variance), changes of replicative age in the differentiated tissue might not always point towards changed dynamics on the stem cell level, but towards abnormal dynamics in the progenitor compartments. Accordingly, we would expect to observe these differences in replicative age distributions in certain diseases that change proliferation and differentiation characteristics in the progenitor compartments. Examples of this include chronic myeloid leukemia, acute promyelocytic leukemia and some other forms of acute myeloid leukemia where a progenitor cell in the ’middle’ of the hierarchy acquires novel properties. For example, increased self renewal would lead to a increase of average cellular age [29, 34, 35, 43].

An important complication that we have not considered here is that real tissues are often found in dynamical regimes that change the cellular age distribution over time. In multicellular organisms the rates for self-renewal and for symmetric differentiation or cell death are variable and tightly regulated by a variety of feedback mechanisms [44]. In this way, a tissue can respond to environmental conditions such as injury or infections. However, we only considered the case of homeostasis and assume that the compartment structure is approximately in a steady state. While this strong assumption allows for a considerable spectrum of observable age distributions in the differentiated tissue, we expect to vary this distribution even more in dynamical settings.

In conclusion, the distribution of replicative age can reveal properties of compartmentalised tissue in multicellular organisms and can inform about changes from healthy tissue due to diseases that change the proliferation characteristics of the cells. Understanding the replicative age distributions of tissue in multicellular organisms can therefore lead to further knowledge of tissue dynamics and ultimately reveal insights about additional disease risks and disease characteristics.

### Availability of source code

The data for the figures in this manuscript was either calculated analytically or solved numerically by using the Scipy library for python. The scripts to create our results figures can be accessed at https://github.com/marvinboe/DownstreamReplAge.

## A. Steady state distribution

Here we show the general solution for the steady state distribution of replicative age inside a downstream compartment for any input distribution 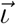. The differential equations for the number of cells in each replicative age class are:

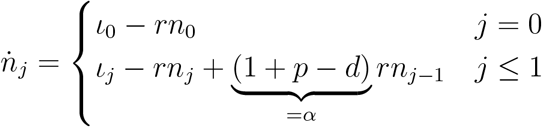

We then start by setting the equation for *n*_0_ to zero, such that

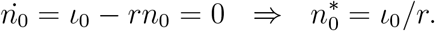

We use this result to solve for *n*_1_ and then continue recursively until we find the general solution for *n*_*j*_:

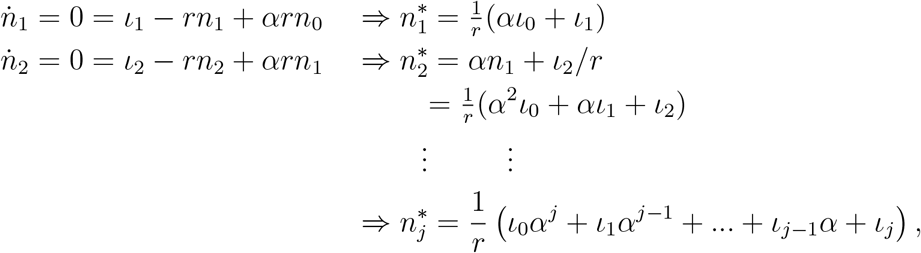

which can be written in the more compact form

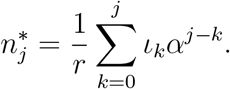

## B. Total cell number amplification

Here we start again by writing down the differential equations for the total number of cells *N*^(*c*)^ in each of the compartments *c* with proliferation rates *r*^(*c*)^. Additionally we have the total influx I into the first compartment (*c* = 0), and similarly the total outflux Ω from the last compartment (*c* = *C* − 1).

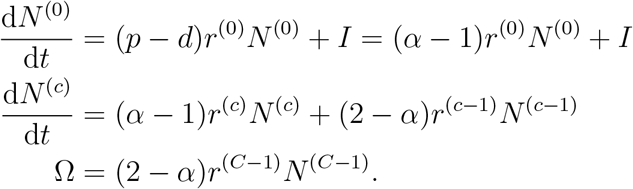

As in the calculation above we assume our compartment to be in the steady state and set the above differential equations to zero.

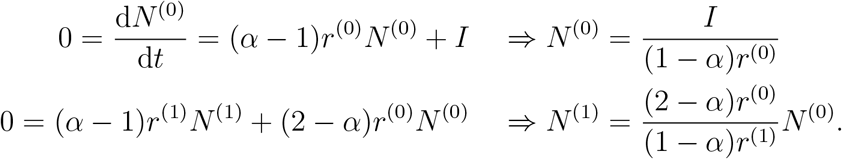

The same calculation can be done for each compartment *c*:

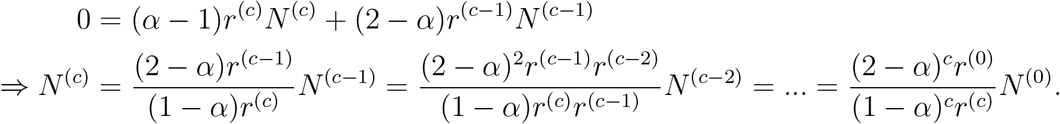

From this follows for the total outflux Ω

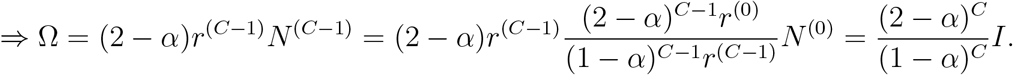

By rearranging this for *α* we get

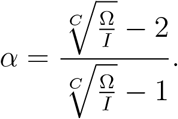

## C. Mean and variance of replicative age distribution

To calculate the moments of the replicative age *j* in the progenitor compartment, we need to normalize the replicative age distribution 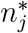 in the steady state by the total number of cells in the progenitor compartment 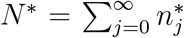. We can then write down the *m*-th moment of the age *j* in the progenitor compartment

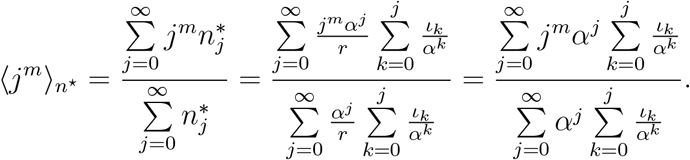

By changing the order of summation in the denominator we get

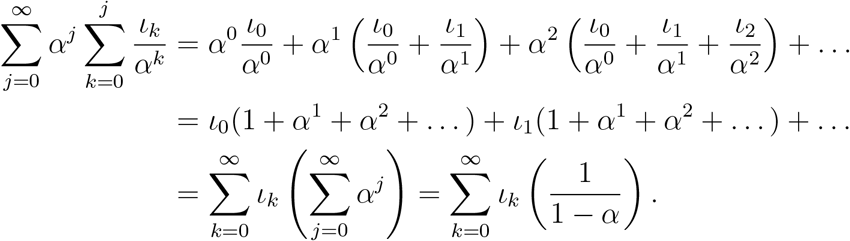

Similarly, we change the order of summation in the nominator to

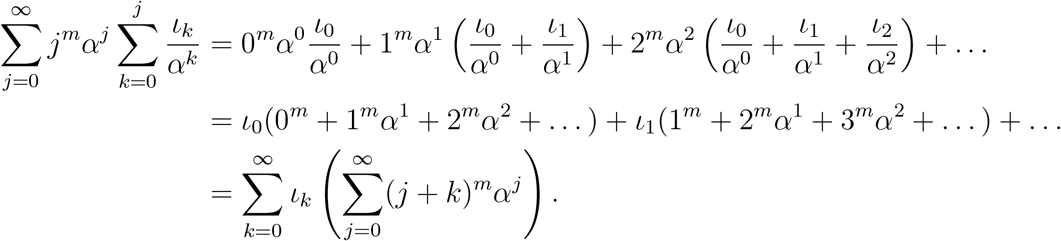

We then rewrite the above binomial to 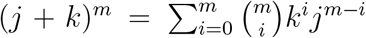 and rearrange the sums to

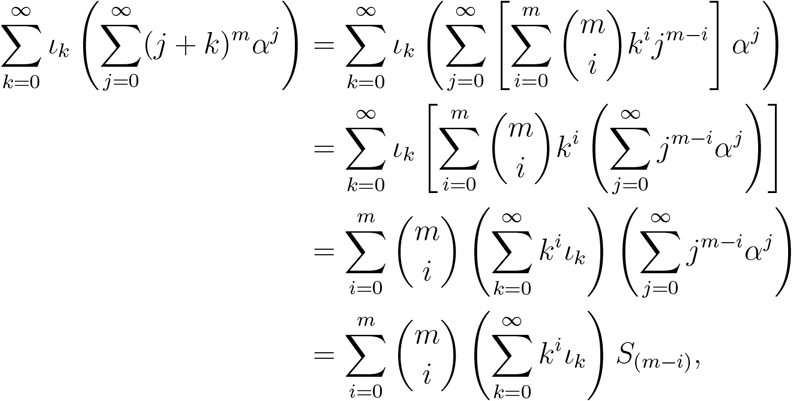

where in the last step we defined the sum which is independent of the influx

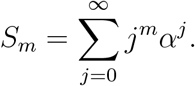

The next step is to put together the nominator and denominator and to insert the moments of the replicative age distribution of the influx 〈*j^m^*〉_*ı*_:

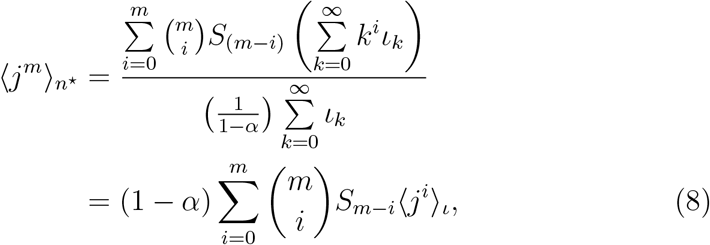

which is the general solution for any moment of the replicative age distribution.

To get an expression for the mean and variance, we have to solve *S_m_* for *m* = 0, 1, 2:

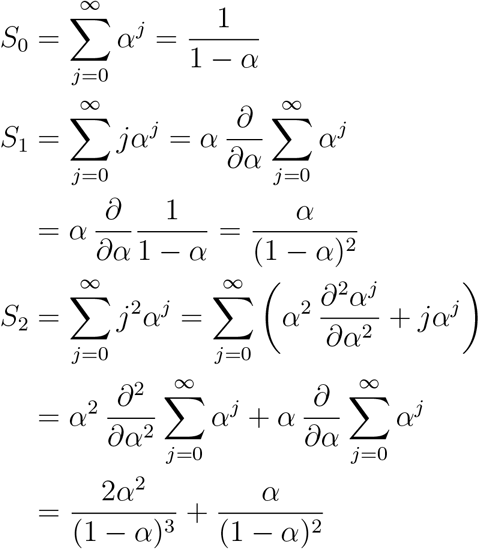

Generally, for *m* > 0 the sum *S*_*m*_ is by definition the polylogarithm *Li_−_*_*m*_(*α*) [45] with negative order *m* and can be written as

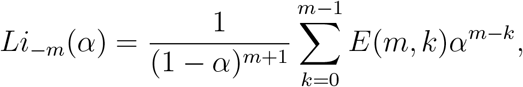

with the Eulerian numbers 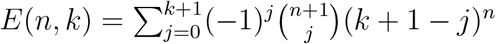.

By using the general solution equation 8 and the above solutions for *S_n_* for *n* = 0, 1, 2 we can calculate the mean *μ* and variance *σ*^2^ of the replicative age distribution:

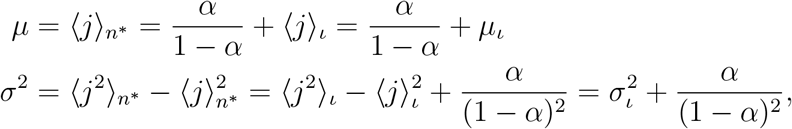

where we used the mean *μ*_*ι*_ and the variance 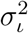 of the influx distribution.

## D. Replicative age distributions for specific influx

### D.1. Geometric influx

Here we calculate the distribution of replicative age in the steady state resulting from geometrically distributed age of the influx *ι_k_* = *λ*^*k*^(1 − *λ*) by solving equation 2:

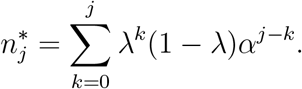

Now we factor out all factors independent of *k* and substitute 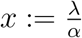:

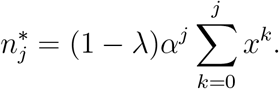

By using the results for the geometric sum 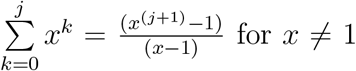 we get

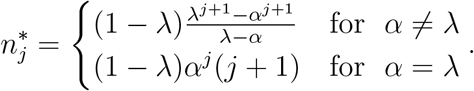

### D.2. Poisson influx

To find the replicative age distribution in the progenitor compartment for Poisson distributed influx, we have to solve the following sum:

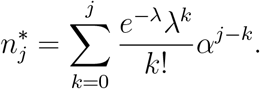

By factoring out all factors independent of *k*, we arrive at

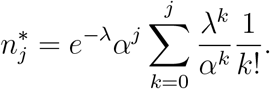

Now we substitute 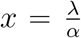 and note that for the upper incomplete gamma function 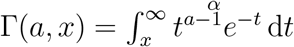 and integer values of *j* the following equality holds [32]:

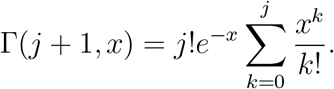

Using this we get

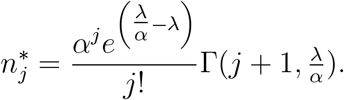

as the desired result.

## E. Time to steady state

Equation 1 can be solved analytically to get an estimate for the time until steady state is reached. In the following we solve the equations for an initially empty progenitor compartment, which gives us an estimate to the relaxation time until steady state is reached. For this we start by solving the equation for the first replicative age class *j* = 0 and then subsequently for all others:

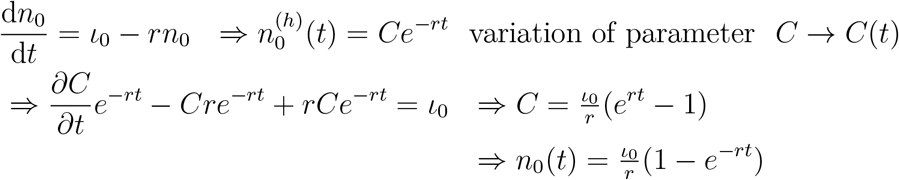

This we plug into the differential equation for the next age class

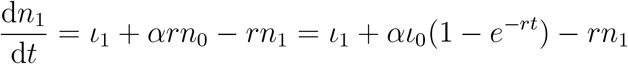

which can be solved in the same way

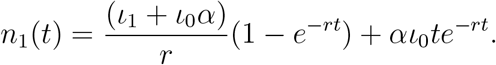

Now we use the steady state values of the main text (equation 2), 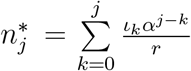, which allows us to rewrite the above equation to

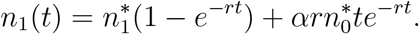

For the third age class we get

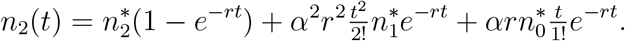

From this we can infer the general solution

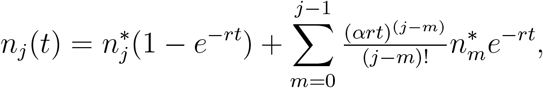

which as expected goes towards the equilibrium solutions 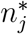 for the limit of time *t* to infinity. Figure E.1A shows the time evolution of of *n_j_* for different age classes *j*. Approach to steady state for this specific value of *α* is relatively fast. To compare the time until steady state is reached for different values of *α*, we numerically calculated the time until *n_j_* reached 99% of the equilibrium value 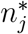 and the results are shown in figure E.1B. As expected, for large *α* the time to reach steady state increases drastically, since many more downstream compartments have to be filled due to the long tail towards old ages in the distribution of replicative age. Also for higher age classes, in this case *j* = 20 for example, time to steady state is much longer as all changes have to go through the previous age classes first. However, since the steady state value in this case is close to zero and the influx into this compartment is even smaller, our assumption of a quasi-static process is still valid.

**Figure E.1:**
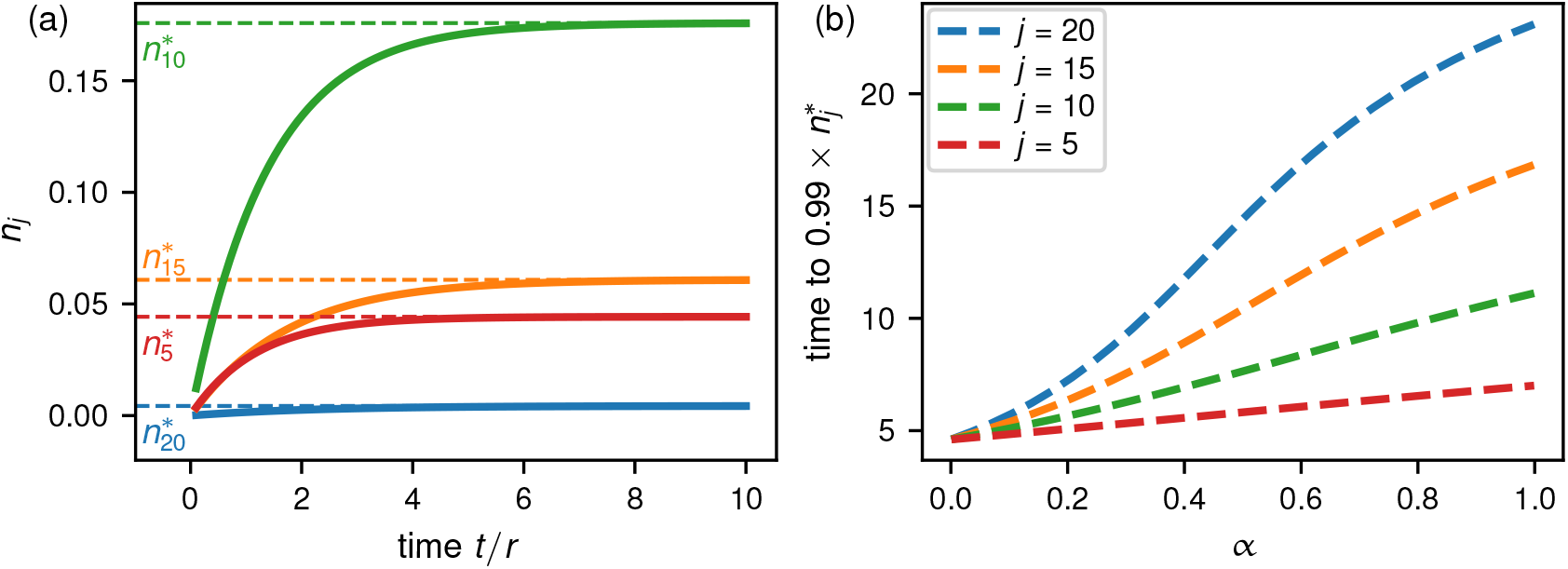
Time to the steady state for different parameters. (a) The number of cells *n_j_* for various age classes *j* increases until the steady state is reached, earlier compartments reach the steady state faster (*α* = 0.3). (b) Time to approach equilibrium solutions for different age classes *j* in a single progenitor compartment for different self-renewal parameters *α*. The influx age is Poisson distributed with *λ* = 10 (see section 3.1.3).

